# *In silico* prediction and characterisation of secondary metabolite clusters in the plant pathogenic fungus *Verticillium dahliae*

**DOI:** 10.1101/481648

**Authors:** Xiaoqian Shi-Kunne, Roger de Pedro Jové, Jasper R.L. Depotter, Malaika Ebert, Michael F. Seidl, Bart P.H.J. Thomma

## Abstract

Fungi are renowned producers of natural compounds, also known as secondary metabolites (SMs) that display a wide array of biological activities. Typically, the genes that are involved in the biosynthesis of SMs are located in close proximity to each other in so-called secondary metabolite clusters (SMCs). Many plant-pathogenic fungi secrete SMs during infection in order to promote disease establishment, for instance as cytocoxic compounds. *Verticillium dahliae* is a notorious plant pathogen that can infect over 200 host plants worldwide. However, the SM repertoire of this vascular pathogen remains mostly uncharted. To unravel the SM potential of *V. dahliae*, we performed *in silico* predictions and in-depth analyses of its SM clusters (SMC). We identified 25 potential SMCs in the *V. dahliae* genome, including loci that can be implicated in DHN-melanin, ferricrocin, triacetyl fusarinine and fujikurin production.

## Introduction

Filamentous fungi are known for their ability to produce a vast array of distinct chemical compounds that are also known as secondary metabolites (SMs) (Keller et al., 2005). In contrast to primary metabolites, SMs are often considered as non-essential for fungal growth, development or reproduction. However, SMs can be crucial for long-term survival in competitive fungal niches (Fox and Howlett, 2008; Ponts, 2015; Derntl et al., 2017). SMs produced by plant pathogenic fungi are of particular interest as they may contribute to virulence, leading to crop losses and threatening food security (Ponts, 2015; Pusztahelyi et al., 2015). For example, T-toxin from *Cochliobolus heterostrophus*, a maize pathogen that caused the worst epidemic in U.S. agricultural history, has been reported to be a crucial pathogenicity factor (Inderbitzin et al., 2010).

Fungal SMs are classified into four main groups based on core enzymes and precursors involved in their biosynthesis: polyketides, non-ribosomal peptides, terpenes and indole alkaloids (Keller et al., 2005). Production of the chemical scaffold of each class requires core enzymes named polyketides synthases (PKSs), non-ribosomal peptide synthetases (NRPSs), terpene cyclases and dimethylallyl tryptophan synthases, respectively. Additionally, hybrid enzymes such as PKS-NRPSs have been identified as builders of structurally complex molecules with combined properties (Boettger and Hertweck, 2013). PKSs and NRPSs are the most abundant and are extensively studied in fungi (Cox, 2007). PKSs can be further divided into three different types (I, II and III), of which type I PKSs and type III PKSs are found in fungi. Type I PKSs are predominant in fungi whereas type III PKSs are found only rarely (Cox, 2007; Gallo et al., 2013; Hashimoto et al., 2014). Genes involved in the synthesis of SMs are frequently located in close proximity to each other, forming so-called secondary metabolite clusters (SMCs) (Keller and Hohn, 1997; Brakhage and Schroeckh, 2011; Wiemann and Keller, 2014). Most of these SMCs contain one biosynthetic core gene that is flanked by transporter proteins, transcription factors, and genes encoding tailoring enzymes that modify the SM structure (Keller & Hohn, 1997; Keller *et al.*, 2005).

The genomics era has provided new tools to study fungal SMs and their biosynthesis at the whole genome scale (Wiemann and Keller, 2014; Medema and Fischbach, 2015). The distinctive traits of gene clusters (e.g. gene distance) and the conserved signatures of core genes (e.g. conserved domains) can be exploited to identify putative loci involved in SM production. Moreover, phylogenetic and comparative genomics analyses are very informative as the number of fungal genomes and characterized SM pathways increases. These two approaches are very helpful to identify gene clusters that are involved in the production of SMs that have been characterized in other fungal species and allow subsequent predictions of identical or related compounds that a particular fungal species might produce (Medema et al., 2013; Cairns and Meyer, 2017).

The fungal genus *Verticillium* contains nine haploid species plus the allodiploid *V. longisporum* (Inderbitzin et al., 2011; Jasper Depotter, Xiaoqian Shi-Kunne, Helene Missionnier, Tingli Liu, Luigi Faino, Grardy van den Berg, Thomas Wood, Baolong Zhang, Alban Jacques, Michael Seidl, 2017). These ten species are phylogenetically subdivided into two clades; Flavexudans and Flavnonexudans (Inderbitzin et al., 2011; Shi-Kunne et al., 2018). The Flavnonexudans clade comprises *V. nubilum, V. alfalfae, V. nonalfalfae, V. dahliae,* and *V. longisporum,* while the Flavexudans clade comprises *V. albo-atrum, V. isaacii, V. tricorpus, V. klebahnii* and *V. zaregamsianum* (Inderbitzin et al., 2011). Among these *Verticillium* spp., *V. dahliae* is the most notorious plant pathogen that is able to cause disease in hundreds of plant species (Fradin and Thomma, 2006; Inderbitzin and Subbarao, 2014). Furthermore, *V. albo-atrum, V. alfalfae, V. nonalfalfae* and *V. longisporum* are pathogenic, albeit with narrower host ranges (Inderbitzin and Subbarao, 2014). Although the remaining species *V. tricorpus, V. zaregamsianum, V. nubilum, V. isaacii* and *V. klebahnii* have incidentally been reported as plant pathogens, they are mostly considered as saprophytes that thrive on dead organic material and their incidental infections should be seen as opportunistic (Ebihara et al., 2003; Inderbitzin et al., 2011; Gurung et al., 2015). Previously, studies of three genes that are involved in SM biosynthesis in *V. dahliae* suggested that SMs may play a role in *V. dahliae* virulence. The deletion mutants of the putative secondary metabolism regulators *VdSge1* (Santhanam and Thomma, 2012) and *VdMcm1* (Xiong et al., 2016) displayed reduced virulence when compared with the wild type *V. dahliae* strain. Likewise, a reduction in virulence was observed for deletion mutants of the cytochrome P450 monooxygenase *VdCYP1* (Zhang et al., 2016), a common tailoring enzyme in SMC production. In this study, we conducted an *in silico* analysis to unravel the potential secondary metabolism of *V. dahliae* by making use of the gapless genome assembly of strain JR2 (Faino et al., 2015)

## RESULTS AND DICUSSION

### The *V. dahliae* strain JR2 genome contains 25 putative secondary metabolite clusters

To assess the potential secondary metabolism of *V. dahliae* strain JR2, we mined its genome sequence to predict SMCs using antiSMASH (Weber et al., 2015). A total of 25 putative SMCs were predicted, containing a total of 364 genes within their boundaries (**Error! Reference source not found. & Error! Reference source not found.**). The putative SMCs were classified as nine type I PKSs, one type III PKS, one PKS-NRPS, three NRPSs and four terpenes. Seven clusters were classified as “other”, a generic class of SMCs containing core enzymes with unusual domain architecture, also known as non-canonical. We found that each of these SMCs contains one core gene. Disrupted SMCs frequently occur in the genomes of filamentous fungi (Collemare et al., 2014). However, none of the clusters identified in *V. dahliae* strain JR2 is evidently disrupted, as all identified clusters have no insertions of transposible elements and include genes encoding tailoring enzymes such as methyltransferases, cytochrome P450 or dehydrogenases (Cacho et al., 2015). Several clusters also comprise transporter and transcription factor encoding genes that might be involved in SM secretion and local gene cluster regulation respectively (Collemare et al., 2014) (Table 1). Collectively, these results suggest that all the analysed SMCs of *V. dahliae* strain JR2 are potentially functional.

In several species, secondary metabolite genes are enriched at chromosomal ends adjacent to the telomeres (Cairns & Meyer, 2017; Farman, 2007; McDonagh et al., 2008). Thus, we assessed whether the SMCs of *V. dahliae* are located in sub-telomeric regions, here defined as within 300 kb from the chromosomal end. We found that 36% of the predicted clusters are in sub-telomeric regions (Figure 1 & Table 1). As these sub-telomeric regions harbour 1606 genes in total, which is 13.8% of total gene repertoire (Table 1) they are significantly enriched (χ2-test, p<0.0001) in secondary metabolism genes (Figure 1).

**Figure 1.**
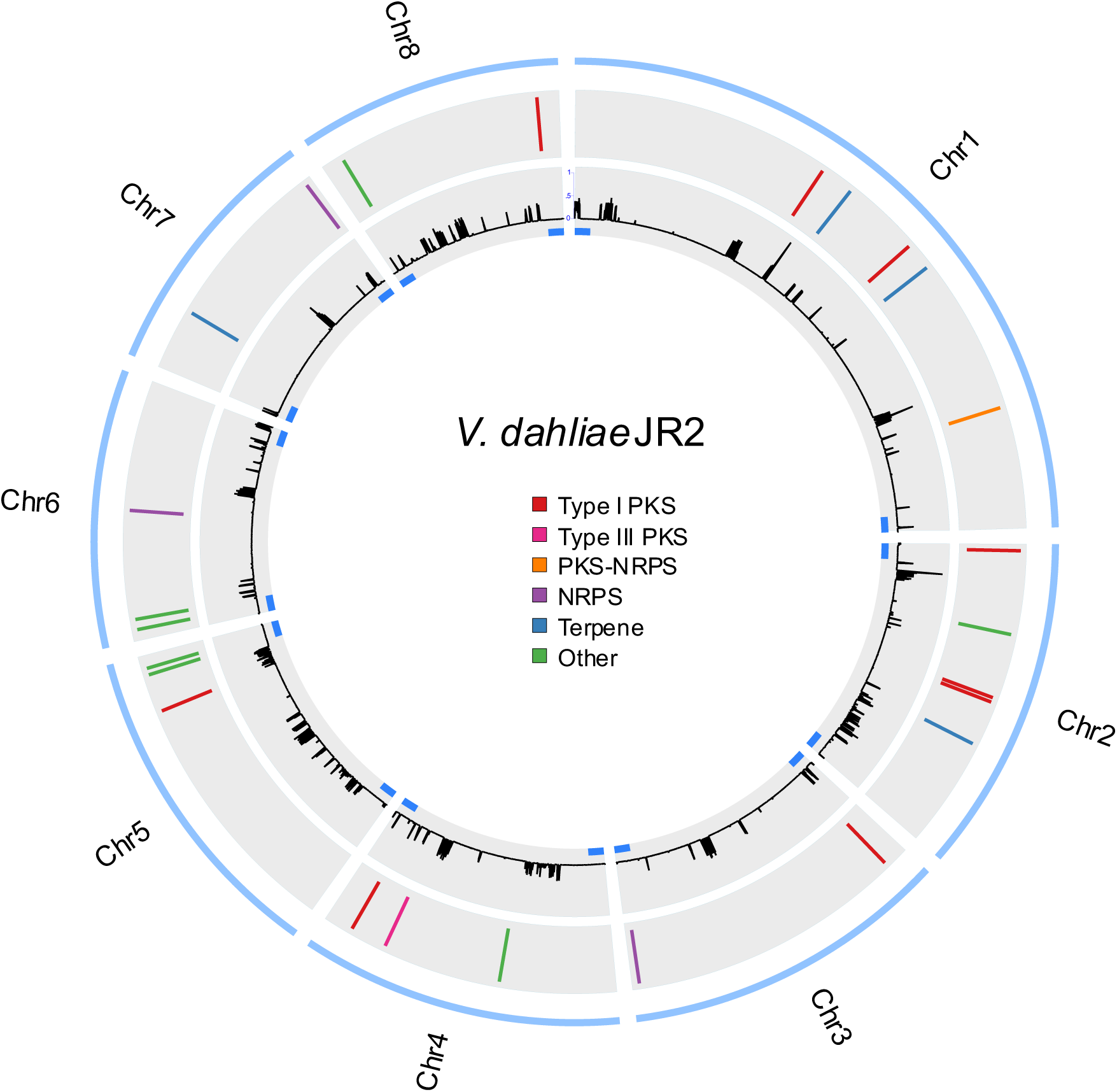
Genomic location of *V. dahliae* SMCs. The outer blue lane represents the chromosomes. The middle grey lane shows the relative position of the predicted SMCs on each chromosome. The inner grey lane shows the repeat density in the JR2 genome. The blue rectangles indicate the regions that are defined as sub-telomeric (300 kb from each chromosomal end).

To check whether the SMCs identified in *V. dahliae* strain JR2 are also present in other *V. dahliae* strains, we assessed the presence/absence of the core enzymes in 22 *V. dahliae* strains (Klosterman et al., 2011; Faino et al., 2015; Kombrink et al., 2017; Depotter et al., 2018). Among these 22 strains is the gapless genome assembly of strain VdLs17 (Klosterman et al., 2011; Faino et al., 2015) and the nearly complete genome assemblies of strains CQ2 and 85S (Depotter et al., 2018). The remaining genome assemblies are considerably fragmented, with over 500 contigs for each of the assemblies. Nevertheless, we found that each of the core enzymes is present in all 22 strians, except for VdPks8 which was next to JR2 only found in VdLs17, CQ2 and 85S. However, the absence of VdPks8 from the other strains may be due to the fragmented genome assemblies. Subsequently, we assessed the genome assembles of strains VdLs17, CQ2 and 85S for the presence of complete clusters, revealing that all SMCs identified in *V. dahliae* strain JR2 are also found in the genomes of these three strians, therefore suggesting that SMCs are highly conserved in *V. dahliae* strains.

To examine whether the SMCs identified in *V. dahliae* strain JR2 are also present in other *Verticillium* spp., we queried core enzymes of each cluster using BLAST against the proteomes of previously published *Verticillium* spp. (Jasper Depotter, Xiaoqian Shi-Kunne, Helene Missionnier, Tingli Liu, Luigi Faino, Grardy van den Berg, Thomas Wood, Baolong Zhang, Alban Jacques, Michael Seidl, 2017; Shi-Kunne et al., 2018). In total, 10 SMC core enzymes are present in all *Verticillium* spp., seven of which showed considerable sequnence conservation (>80% sequence identity) (Figure 2). The other 15 core enzymes are not present in all species, but display a mosaic presence/absence pattern with presence in at least two other species. Of these, VdPks7 is conserved in all species except for *V. longisporum* (Figure 2). VdPks3 is only conserved in the closely related species *V. alfalfae* and *V. nonalfalfae* and VdPks5 is conserved in all pathogenic species except *V. albo-atrum*. Interestingly, the VdPks8 core enzyme was only found in a single copy in the hybrid *V. longisporum* genome, presumably derived from its *V. dahliae* progenitor. Thus, based on the widespread presence of these core genes within the *Verticillium* genus, we predict that most of the SMCs are conserved throughout this genus.

**Figure 2.**
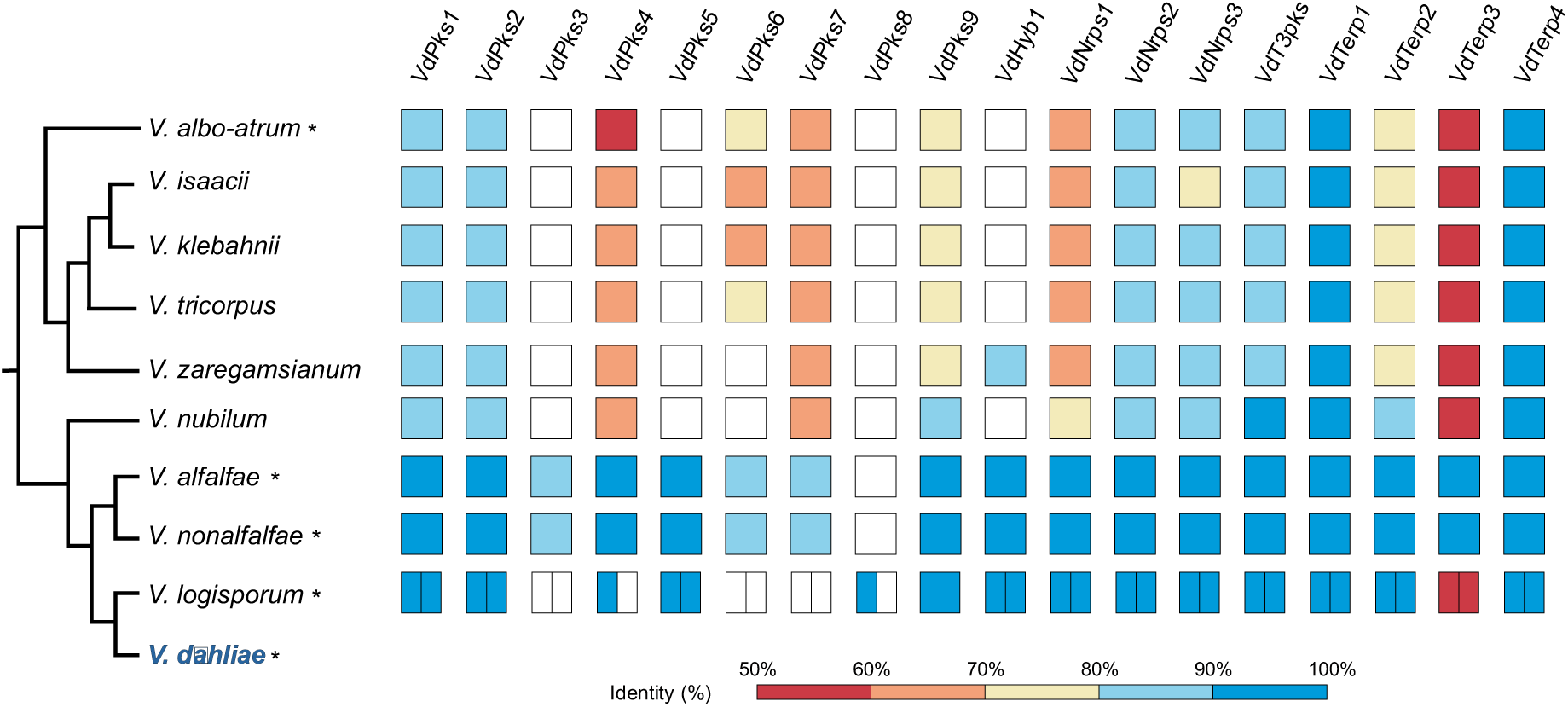
Conservation of *Verticillium dahliae* core SMC genes throughout the *Verticillium* genus. The colour gradient represents the % identity range of the high scoring. Species described as plant pathogens are indicated with an asterisk.

### Phylogenomic analysis of *V. dahliae* secondary metabolite core enzymes

In ascomycetes, type I PKSs, NRPSs and PKS-NRPSs are known to produce most of the SMs that are involved in virulence (Pusztahelyi et al., 2015). Thus, we focused on identifying putative functions of type I PKSs, NRPSs and PKS-NRPSs in *V. dahliae* with a phylogenomics approch. To this end, we aligned KS domians of *V. dahliae* PKSs to KS domains of functionally described PKSs, and subsequently constructed a phylogenetic tree that comprises three major clades that correspond to the NR-PKSs, PR-PKSs and HR-PKSs, respectively (Figure 3). The NR-PKS clade contains two predicted *V. dahliae* PKSs, VdPks2 and VdPks3. VdPks2 clusters with PKSs that have been implicated in dyhydroxynaphthalene (DHN)-melanin formation (Tsuji et al., 2002; Yu et al., 2015). VdPks3 clustered with the Orsellinic acid synthase OpS1, which is involved in production of the toxic metabolite oosporein by the entomopathogenic fungus *Beauveria bassiana* (Feng et al., 2015). The HR-PKS clade showed that four out of the seven predicted *V. dahliae* HR-PKSs (VdPks1, VdPks7, VdPks8 and VdPks9) grouped with previously described enzymes. VdPks7 and VdPks8 clustered with the fumagillin synthase from *Aspergillus fumigatus* and fujikurin synthase from *Fusarium fujikuroi*, respectively (Lin et al., 2013; von Bargen et al., 2015; Niehaus et al., 2016). VdPks1 and VdPks9 clustered with the T-toxin synthase PKS2 from *Cochliobolus heterostrophus* (Inderbitzin et al., 2010). VdPks4 grouped in a clade that contians sdnO and PKS1 synthase, which are involved in the production of Sordarin by *Sordaria araneosa* (Kudo et al., 2016), an antifungal agent that inhibits protein synthesis in fungi by stabilizing the ribosome/EF2 complex (Justice et al., 1998), and in T-toxin production in *C. heterostrophus* (Inderbitzin et al., 2010), respectively. VdPks5 is in a clade that only contains FUB1 fusaric acid synthase orthologs of three *Fusarium* spp. (Brown et al., 2015). The remaining *V. dahliae* PKS core enzyme, VdPks6, is not directly grouping adjacent to any previously described enzyme (Figure 3). Thus, eight of the nine *V. dahliae* PKS core enzymes (VdPks1, VdPks2, VdPks3, VdPks4, VdPks5, VdPks7, VdPks8, VdPks9 and VdHyb1) group with previously characterised enzymes, thereby allowing us to infer their putative function.

**Figure 3.**
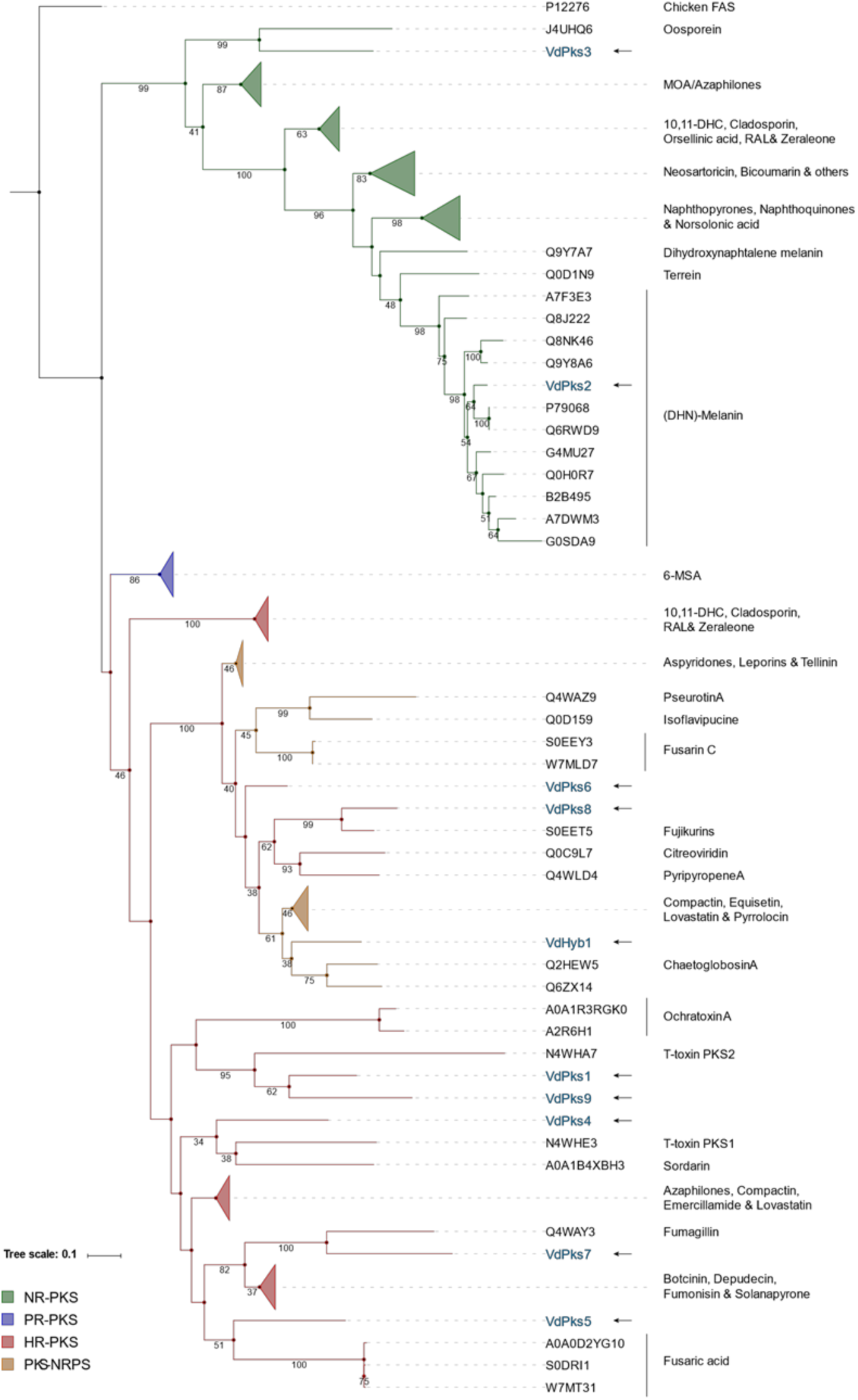
Phylogenetic tree of type I PKS and PKS-NRPS enzymes. KS domains of PKS and PKS-NRPS enzymes were aligned to construct the maximum likelihood tree with 100 bootstrap replicates. The chicken fatty acid synthase (Chicken FAS) sequence was used as outgroup. Only bootstrap values above 30 are shown below the branches. *V. dahliae* KS domains are highlighted in blue and indicated with an arrow. Protein codes correspond to Uniprot IDs.

Like for PKSs, we similarly perfomed phylogenomic analysis to get more insight into the putative products of NRPSs. The conserved A-domain sequences of *V. dahliae* NRPS core enzymes were aligned with previously described enzymes of other fungal species to construct a phylogenetic tree. A distinct clade clearly separated NRPSs from the PKS-NRPSs in the phylogenetic tree (Figure 4). VdNprs1 grouped in a clade with NPS2, which is involved in the synthesis of the intracellular siderophore ferricrocin of *Fusarium pseudograminearum* and *Cochliobolus heterostrophus* (Tobiasen et al., 2007; Sieber et al., 2014; Oide et al., 2015). VdNprs2 clusters with NRPS4, which is responsible for the synthesis of the extracellular siderophore triacetylfusarinine C (TAFC) by *A. fumigatus* (Schrettl et al., 2007). The clade that contains VdNrps3 has low bootstrap values and long branches, indicating considerable divergence of this enzyme (Figure 4). Thus, only two of the *V. dahliae* NRPS core enzymes (VdNprs1 and VdNprs2) group with previously characterised enzymes.

**Figure 4.**
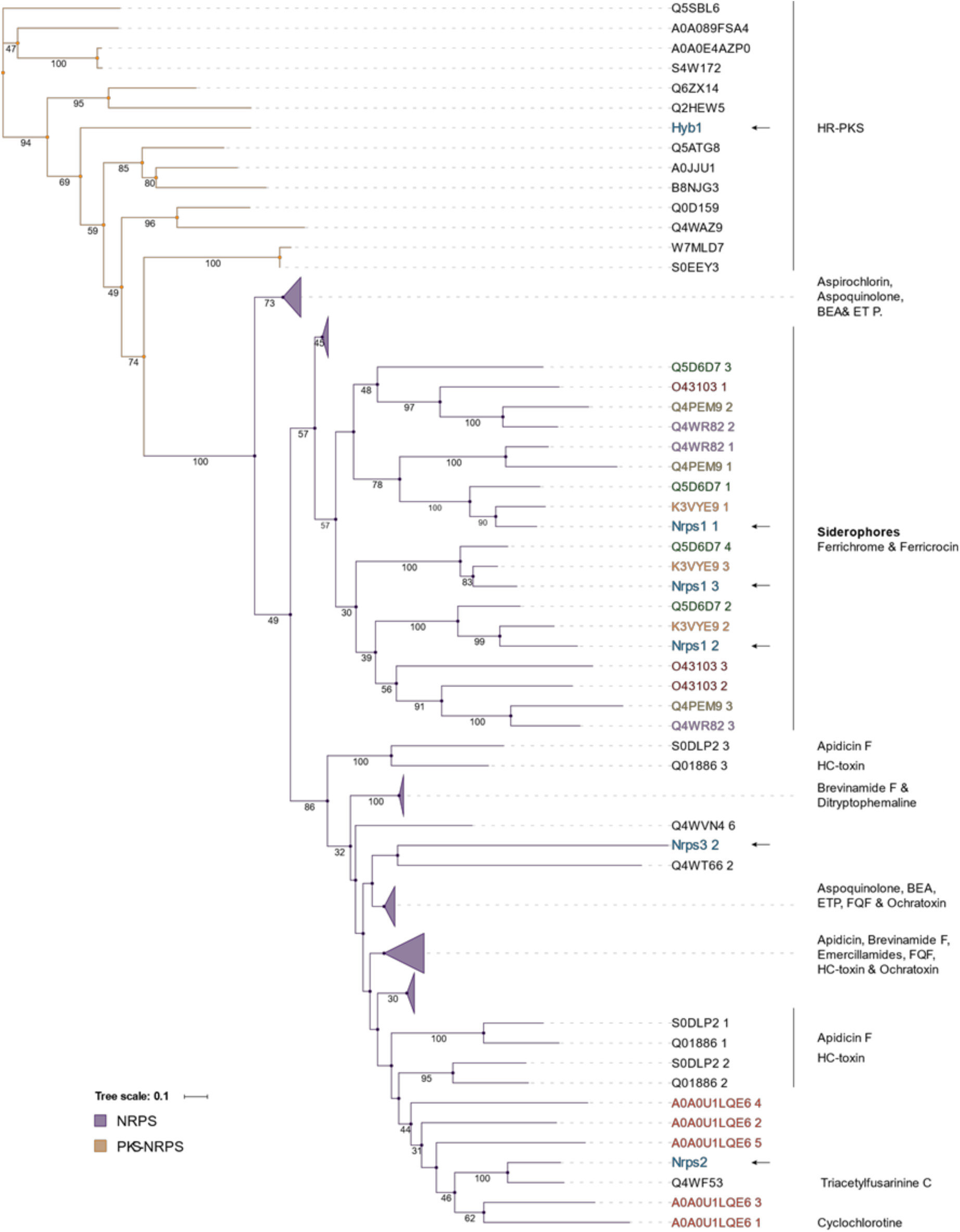
Phylogenetic tree of NRPS and PKS-NRPS enzymes. A-domains of NRPS and PKS-NRPS enzymes were aligned to construct the maximum likelihood tree with 100 bootstrap replicates. Only bootstrap values over 30 are shown below the branches. *V. dahliae* A-domains are highlighted in blue and indicated with an arrow.

The fusion of PKS and NRPS domains results in PKS-NRPS enzymes that stand out due to their structural complexity (Boettger and Hertweck, 2013). In the genome of *V. dahliae* (strain JR2), only one PKS-NRPS (VdHyb1) was detected. Since it contained a KS domain characteristic for HR-PKSs and an A-domain as commonly observed in NRPSs, we included VdHyb1 in both phylogenetic trees. In the PKS phylogeny (Figure 3) VdHyb1 was found in the same clade (38%, bootstrap value) as the PKS-NRPSs chaetoglobosin A synthase from *Chaetomium globosum* and the avirulence protein ACE1 from *Magnaporthe grisea* (Collemare et al., 2008; Ishiuchi et al., 2013). In contrast, VdHyb1 did not cluster with any previously characterized PKS-NRPS in the A-domain phylogenetic tree (Figure 4).

### Comparative analysis of gene clusters

Based on the phylogenetic analyses, 11 core enzymes (VdPks1, VdPks2, VdPks3, VdPks4, VdPks5, VdPks7, VdPks8, VdPks9, VdNrps1, VdNrps2 and VdHyb1) were identified that cluster with previously characterised enzymes from other fungal species (Figure 3 & Figure 4). Subsequently, we queried for the other genes besides the core genes from these previously characterised clusters in other fungal species to find homologs in the corresponding *V. dahliae* clusters. However, only the VdPks2, VdPks8, VdNrps1 and VdNrps2 clusters of *V. dahliae* share more homologs (more than 50% of the whole cluster) in addition to the core genes with other fungal species. The remaining clusters contain less than 50% of genes that share homologs with other fungal species. In other fungi, conserved gene clusters of VdPks2, VdPks8, VdNrps1 and VdNrps2 are responsible for the biosynthesis of DHN-melanin, fujikurins, ferricrocin and TAFC, respectively (Tsuji et al., 2002; Tobiasen et al., 2007; Sieber et al., 2014; von Bargen et al., 2015; Niehaus et al., 2016).

The gene cluster for DHN-melanin biosynthesis in the fungal pathogen *C. lagenarium* comprises six genes, including three functionally characterized genes encoding polyketide synthase *ClPKS1*, reductase *T4HR1* and transcription factor *cmr1*. Moreover, two genes (scytalone dehydratase *SCD1* and *THR1* reductase) residing at another chromosome were identified to be involved in the biosynthesis of DHN-melanin as well (Tsuji et al., 2002) (Figure 5A). We observed amino acid identities of 70%-90% between ClPKS1, T4HR1, SCD1 and THR1 of *C. lagenarium* and their orthologs in *V. dahliae*. The transcription factor CMR1 only shares 55% identity with its counterpart in *C. lagenarium*.

**Figure 5.**
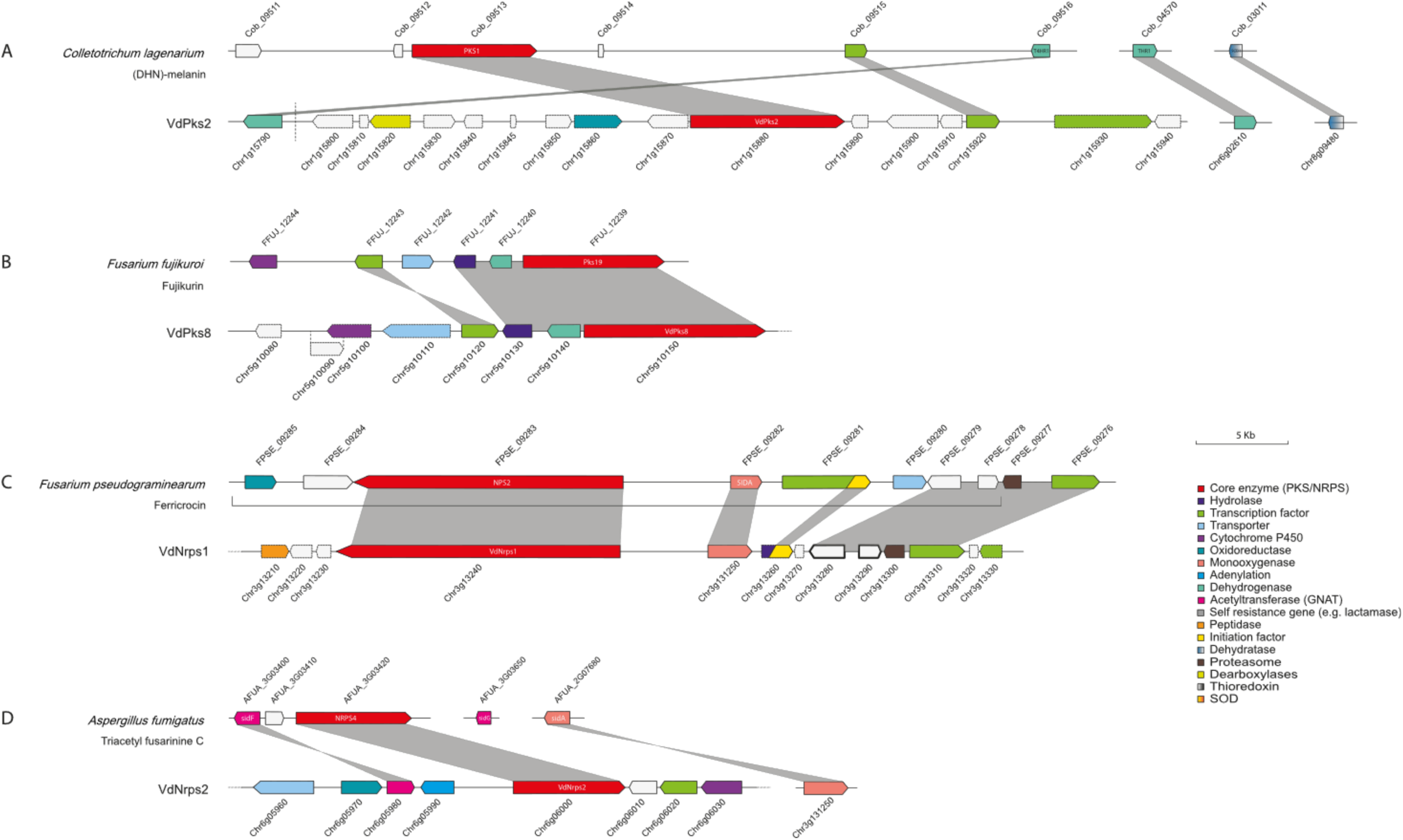
Synteny of conserved SMCs in *V. dahliae*. *V. dahliae* putative clusters were compared to previously described clusters. Ensembl gene IDs are shown above or below the genes. (A) DHN-melanin, (B) Fujikurins, (C) Ferricrocin, (D) Triacetyl fusarinine

*VdPks8* is an ortholog of the fujikurin synthase gene (*FfuPks19*) of *F. fujikuroi*. The fujikurin cluster contains six genes (von Bargen et al., 2015; Niehaus et al., 2016), four of which have homologs in the *VdPks8* locus in *V. dahliae*. The homolog of the MFS transporter FFUJ_12242 of *F. fujikuroi* was found on a different chromosome in *V. dahliae,* and the cytochrome P450 gene in *F. fujikuroi* has no *V. dahliae* homolog (Figure 5B). Interestingly, the *VdPks8* locus contains two other genes that are annotated as cytochrome P450 and MFS transporter, but these genes were not detected as homologs of the genes in the *FfuPks19* cluster (Figure 5B).

The biosynthesis of ferricrocin requires two genes that are located at the same locus in the genome of *F. pseudograminearum*, which encode an L-ornithine N5-oxygenase (SIDA) and an NRPS (FpNRPS2) (Tobiasen et al., 2007; Sieber et al., 2014). Similarly, in *V. dahliae* homologs of L-ornithine N5-oxygenase (SIDA) and NRPS (FpNRPS2) genes are located next to each other. Moreover, homologs of a proteasome subunit, a transcription factor and two uncharacterised genes in the same cluster of *F. pseudograminearum* were found in *V. dahliae.* In addition, genes encoding an MFS transporter and an oxireductase in *F. pseudograminearum* have no homologs in *V. dahliae* (Figure 5C).

VdNrps2 is an ortholog of the extracellular siderophore TFAC synthase gene *NRPS4* in *A. fumigatus,* which belongs to a cluster of two genes (*NRPS4* and sidF) (Schrettl et al., 2007). Another two extracellular siderophore TFAC synthase genes (*sidG and sidA*) are located at another chromosome (Schrettl et al., 2007). Except for *sidG*, all described genes have homologs in *V. dahliae* (Figure 5D).

To investigate the putative role of the SM clusters in plant pathogen interactions, we compared expression patterns of each core gene *in planta* and *in vitro*. We assessed transcription (RNA-seq) data of *V. dahliae* during colonization of *Arabidopsis thaliana* and found that all four core genes of the four conserved clusters showed *in planta* expression. Whereas *VdNrps1* was found to be induced, *VdPks2*, *VdPks8*, and *VdNrps2* were repressed when compared with the expression in *in vitro*-cultured mycelium (Figure 6). Nevertheless, the expression *in planta* suggests that these SM cluster may play a role during host colonization.

**Figure 6.**
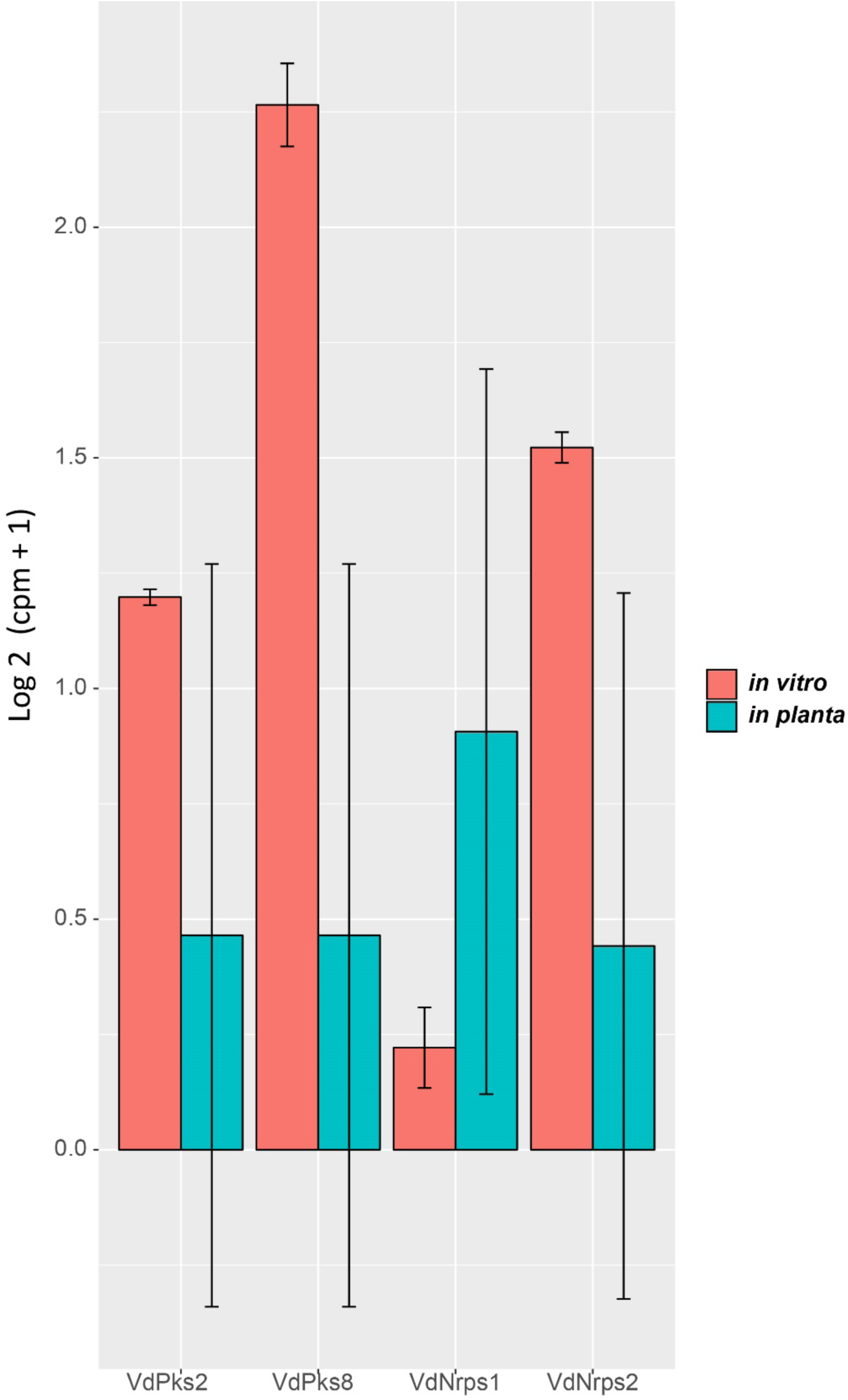
Pair-wise comparison of core SMC genes with differential expression *in vitro* and *in planta*. Gene expressions are depicted for *V. dahliae* strain JR2 cultured in liquid medium and upon *A. thaliana* colonization, respectively. Bars represent the mean gene expression with standard deviations. The significance of difference in gene expression was calculated using t-tests relative to a threshold (TREAT) of log2-fold-change ≥ 1 (McCarthy and Smyth, 2009).

## Conclusions

In this study, we have used an *in silico* approach to identify 25 putative SMCs in the genome of *V. dahliae* strain JR2, all of which appear complete and thus potentially functional. Our predictions state that two putative siderophores, ferricrocin and TAFC, DHN-melanin and fujikurin compounds may belong to the active SM repertoire of *V. dahliae*.

## Material and Methods

### Secondary metabolite cluster prediction, annotation and conservation

Putative SMCs were identified with antiSMASH fungal version 4.0.2 (Weber et al., 2015). The predicted borders from AntiSMASH were directly used to retrieve all protein sequences contained within the clusters. The Bedtools intersect command (Quinlan & Hall, 2010) was used to obtain the file containing the gene locations, followed by gffread from the Cufflinks package (Trapnell et al., 2010) to retrieve the protein sequences. Sub-telomeric regions were defined as 300 kb of the chromosomal ends, as similarly used for other filamentous fungi (McDonagh et al., 2008; Cairns and Meyer, 2017). Genes within the genomic range were counted using BioMart from Ensembl (Kersey et al., 2016). A χ^2^ test was performed to determine the significance of enrichment.

The conservation of predicted SMCs among *Verticillium* spp. was assessed based on core enzyme conservation, using BLAST+ tool protein blast (blastp) (Camacho et al., 2009) on predicted protein databases (e-value < 1 × 10^-5^, query coverage > 60 % and identity > 50 % (Sbaraini et al., 2017).

### Phylogenetic analysis

The previously described type I PKSs, NRPSs and PKS-NRPSs enzymes used in this study for phylogenetic analysis were derived from the curated database of UniProt, SwissProt and literatures (Gallo et al., 2013; Yu et al., 2015). The amino acid alignment was built using MAFFT version 7.205 (Katoh and Standley, 2013). We used the G-INS-i strategy, global alignment (--globalpair) and 1000 cycles of iterative refinement (--maxiterate 1000). Aligned sequences were visualised with Aliview version 1.20 (Larsson, 2014) and manually curated by removing non-aligned sequences. Preceding the phylogenetic analysis, the alignments were trimmed to remove poorly aligned regions using TrimAl version 1.4 (Capella-Gutiérrez et al., 2009). First, all positions in the alignment with gaps in 90% or more of the sequence were removed (-gt 0.1), followed by the automated1 parameter (-automated1). RaxML version 8.1.1 (Stamatakis, 2014) was used to construct Maximum-likelihood phylogenetic tree (-f a). The automated protein model selection (-m PROTGAMMAAUTO) was used applying 100 rapid bootstrapping (-#100). The number of seeds for parsimony inferences and rapid bootstrap analysis was set to 12345 (-p 12345 -x 12345, respectively). The output tree (RaxML_bipartitionBranchLabels) was visualised using iTOL webtool version 3.0 (Letunic and Bork, 2016).

### Comparative cluster analysis

Protein sequences of described clusters were blasted (BLASTp, E-value cutoff 1e-5, query coverage > 60 % and identity > 25 %) against the *V. dahliae* strain JR2 protein database. We considered a cluster to be conserved in *V. dahliae* when at least 50 % of the queried proteins from previously described clusters were found in *V. dahliae*.

### Gene expression analysis

To obtain RNA-seq data for *V. dahliae* grown in culture medium, strain JR2 was grown for three days in potato dextrose broth (PDB) in three biological replicates. To obtain RNA-seq data from *V. dahliae* grown *in planta*, three-week-old *A. thaliana* (Col-0) plants were inoculated with strain JR2. After root inoculation, plants were grown in individual pots in a greenhouse under a cycle of 16 h of light and 8 h of darkness, with temperatures maintained between 20 and 22°C during the day and a minimum of 15°C overnight. Three pooled samples (10 plants per sample) of complete flowering stems were used for total RNA extraction. Total RNA was extracted based on TRIzol RNA extraction (Simms et al., 1993). cDNA synthesis, library preparation (TruSeq RNA-Seq short insert library), and Illumina sequencing (single-end 50 bp) was performed at the Beijing Genome Institute (BGI, Hong Kong, China). In total, ∼2 Gb and ∼1.5 Gb of filtered reads were obtained for the *V. dahliae* samples grown in culture medium and *in planta*, respectively. RNAseq data were submitted to the SRA database under the accession number: SRP149060.

The RNA sequencing reads were mapped to their previously assembled genomes using the Rsubread package in R (Liao et al., 2013). The comparative transcriptomic analysis was performed with the package edgeR in R (v3.4.3) (Robinson et al., 2010; McCarthy et al., 2012). Genes are considered differentially expressed when P-value < 0.05 with a log2-fold-change ≥ 1. P-values were corrected for multiple comparisons according to Benjamini and Hochberg (Benjamini and Hochberg, 1995).

## ACKNOWLEDGEMENTS

This work was supported by the Research Council Earth and Life Sciences (ALW) of the Netherlands Organization of Scientific Research (NWO) to B.P.H.J.T. and M.F.S. The authors declare no conflict of interest.

## REFERENCES

von Bargen KW, Niehaus E-M, Krug I, Bergander K, Würthwein E-U, Tudzynski B, Humpf H-U (2015) Isolation and structure elucidation of fujikurins A–D: products of the PKS19 gene cluster in Fusarium fujikuroi. J Nat Prod 78: 1809–1815

Benjamini Y, Hochberg Y (1995) Controlling the false discovery rate: a practical and powerful approach to multiple testing. J R Stat Soc Ser B 289–300

Boettger D, Hertweck C (2013) Molecular diversity sculpted by fungal PKS–NRPS hybrids. ChemBioChem 14: 28–42

Brakhage AA, Schroeckh V (2011) Fungal secondary metabolites–strategies to activate silent gene clusters. Fungal Genet Biol 48: 15–22

Brown DW, Lee S-H, Kim L-H, Ryu J-G, Lee S, Seo Y, Kim YH, Busman M, Yun S-H, Proctor RH (2015) Identification of a 12-gene fusaric acid biosynthetic gene cluster in Fusarium species through comparative and functional genomics. Mol Plant-Microbe Interact 28: 319–332

Cacho RA, Tang Y, Chooi Y-H (2015) Next-generation sequencing approach for connecting secondary metabolites to biosynthetic gene clusters in fungi. Front Microbiol 5: 774

Cairns T, Meyer V (2017) In silico prediction and characterization of secondary metabolite biosynthetic gene clusters in the wheat pathogen Zymoseptoria tritici. BMC Genomics 18: 631

Camacho C, Coulouris G, Avagyan V, Ma N, Papadopoulos J, Bealer K, Madden TL (2009) BLAST+: architecture and applications. BMC Bioinformatics 10: 421

Capella-Gutiérrez S, Silla-Martínez JM, Gabaldón T (2009) trimAl: a tool for automated alignment trimming in large-scale phylogenetic analyses. Bioinformatics 25: 1972–1973

Collemare J, Griffiths S, Iida Y, Jashni MK, Battaglia E, Cox RJ, de Wit PJGM (2014) Secondary metabolism and biotrophic lifestyle in the tomato pathogen Cladosporium fulvum. PLOS One 9: e85877

Collemare J, Pianfetti M, Houlle A, Morin D, Camborde L, Gagey M, Barbisan C, Fudal I, Lebrun M, Böhnert HU (2008) Magnaporthe grisea avirulence gene ACE1 belongs to an infection-specific gene cluster involved in secondary metabolism. New Phytol 179: 196–208

Cox RJ (2007) Polyketides, proteins and genes in fungi: programmed nano-machines begin to reveal their secrets. Org Bomolecular Chem 5: 2010–2026

Depotter J, Shi-Kunne X, Missionnier H, Liu T, Faino L, van den Berg G, Wood T, Zhang B, Jacques A, Seidl M, et al (2018) Dynamic virulence-related regions of the fungal plant pathogen Verticillium dahliae display remarkably enhanced sequence conservation. bioRxiv 277558

Derntl C, Kluger B, Bueschl C, Schuhmacher R, Mach RL, Mach-Aigner AR (2017) Transcription factor Xpp1 is a switch between primary and secondary fungal metabolism. Proc Natl Acad Sci U S A 114: E560–E569

Ebihara Y, Uematsu S, Nagao H, Moriwaki J, Kimishima E (2003) First report of Verticillium tricorpus isolated from potato tubers in Japan. Mycoscience 44: 481–488

Faino L, Seidl MF, Datema E, van den Berg GCM, Janssen A, Wittenberg AHJ, Thomma BPHJ (2015) Single-molecule real-time sequencing combined with optical mapping yields completely finished fungal genome. MBio 6: e00936–15

Feng P, Shang Y, Cen K, Wang C (2015) Fungal biosynthesis of the bibenzoquinone oosporein to evade insect immunity. Proc Natl Acad Sci U S A 112: 11365–11370

Fox EM, Howlett BJ (2008) Secondary metabolism: regulation and role in fungal biology. Curr Opin Microbiol 11: 481–487

Fradin EF, Thomma BPHJ (2006) Physiology and molecular aspects of Verticillium wilt diseases caused by V. dahliae and V. albo-atrum. Mol Plant Pathol 7: 71–86

Gallo A, Ferrara M, Perrone G (2013) Phylogenetic study of polyketide synthases and nonribosomal peptide synthetases involved in the biosynthesis of mycotoxins. Toxins. doi:10.3390/toxins5040717

Gurung S, Short DPG, Hu X, Sandoya G V, Hayes RJ, Koike ST, Subbarao K V (2015) Host range of Verticillium isaacii and Verticillium klebahnii from artichoke, spinach, and lettuce. Plant Dis 99: 933–938

Hashimoto M, Nonaka T, Fujii I (2014) Fungal type III polyketide synthases. Nat Prod Rep 31: 1306–1317

Inderbitzin P, Asvarak T, Turgeon BG (2010) Six new genes required for production of T-toxin, a polyketide determinant of high virulence of Cochliobolus heterostrophus to Maize. Mol Plant-Microbe Interact 23: 458–472

Inderbitzin P, Bostock RM, Davis RM, Usami T, Platt HW, Subbarao K V (2011) Phylogenetics and taxonomy of the fungal vascular wilt pathogen Verticillium, with the descriptions of five new species. doi:10.1371/journal.pone.0028341

Inderbitzin P, Subbarao K V (2014) Verticillium systematics and evolution: how confusion impedes Verticillium wilt management and how to resolve it. Phytopathology 104: 564–574

Ishiuchi K, Nakazawa T, Yagishita F, Mino T, Noguchi H, Hotta K, Watanabe K (2013) Combinatorial generation of complexity by redox enzymes in the chaetoglobosin A biosynthesis. J Am Chem Soc 135: 7371–7377

Jasper Depotter, Xiaoqian Shi-Kunne, Helene Missionnier, Tingli Liu, Luigi Faino, Grardy van den Berg, Thomas Wood, Baolong Zhang, Alban Jacques, Michael Seidl VOPT (2017) A distinct and genetically diverse lineage of the hybrid fungal pathogen Verticillium longisporum population causes stem striping in British oilseed rape. Environ Microbiol 19: 3997–4009

Justice MC, Hsu M-J, Tse B, Ku T, Balkovec J, Schmatz D, Nielsen J (1998) Elongation factor 2 as a novel target for selective inhibition of fungal protein synthesis. J Biol Chem 273: 3148–3151

Katoh K, Standley DM (2013) MAFFT multiple sequence alignment software version 7: improvements in performance and usability. Mol Biol Evol 30: 772–780

Keller NP, Hohn TM (1997) Metabolic pathway gene clusters in filamentous fungi. Fungal Genet Biol 21: 17–29

Keller NP, Turner G, Bennett JW (2005) Fungal secondary metabolism—from biochemistry to genomics. Nat Rev Microbiol 3: 937

Kersey PJ, Allen JE, Armean I, Boddu S, Bolt BJ, Carvalho-Silva D, Christensen M, Davis P, Falin LJ, Grabmueller C, et al (2016) Ensembl Genomes 2016: more genomes, more complexity. Nucleic Acids Res 44: D574–D580

Klosterman SJ, Subbarao K V., Kang S, Veronese P, Gold SE, Thomma BPHJ, Chen Z, Henrissat B, Lee YH, Park J, et al (2011) Comparative genomics yields insights into niche adaptation of plant vascular wilt pathogens. PLOS Pathog. doi:10.1371/journal.ppat.1002137

Kombrink A, Rovenich H, Shi-kunne X, Rojas-padilla E, van den Berg GCM, Domazakis E, de Jonge R, Valkenburg D-J, Sánchez-Vallet A, Seidl MF, et al (2017) Verticillium dahliae LysM effectors differentially contribute to virulence on plant hosts. Mol Plant Pathol 8: 596–608

Kudo F, Matsuura Y, Hayashi T, Fukushima M, Eguchi T (2016) Genome mining of the sordarin biosynthetic gene cluster from Sordaria araneosa Cain ATCC 36386: characterization of cycloaraneosene synthase and GDP-6-deoxyaltrose transferase. J Antibiot 69: 541

Letunic I, Bork P (2016) Interactive tree of life (iTOL) v3: an online tool for the display and annotation of phylogenetic and other trees. Nucleic Acids Res 44: W242–W245

Liao Y, Smyth GK, Shi W (2013) The Subread aligner: fast, accurate and scalable read mapping by seed-and-vote. Nucleic Acids Res 41: e108–e108

Lin H-C, Chooi Y-H, Dhingra S, Xu W, Calvo AM, Tang Y (2013) The fumagillin biosynthetic gene cluster in Aspergillus fumigatus encodes a cryptic terpene cyclase involved in the formation of β-trans-Bergamotene. J Am Chem Soc 135: 4616–4619

McCarthy DJ, Chen Y, Smyth GK (2012) Differential expression analysis of multifactor RNA-Seq experiments with respect to biological variation. Nucleic Acids Res 40: 4288–4297

McCarthy DJ, Smyth GK (2009) Testing significance relative to a fold-change threshold is a TREAT. Bioinformatics 25: 765–771

McDonagh A, Fedorova ND, Crabtree J, Yu Y, Kim S, Chen D, Loss O, Cairns T, Goldman G, Armstrong-James D, et al (2008) Sub-telomere directed gene expression during initiation of invasive aspergillosis. PLOS Pathog 4: e1000154

Medema MH, Fischbach MA (2015) Computational approaches to natural product discovery. Nat Chem Biol 11: 639

Medema MH, Takano E, Breitling R (2013) Detecting sequence homology at the gene cluster level with MultiGeneBlast. Mol Biol Evol 30: 1218–1223

Niehaus E-M, Münsterkötter M, Proctor RH, Brown DW, Sharon A, Idan Y, Oren-Young L, Sieber CM, Novák O, Pencík A, et al (2016) Comparative “Omics” of the Fusarium fujikuroi species complex highlights differences in genetic potential and metabolite synthesis. Genome Biol Evol 8: 3574–3599

Oide S, Berthiller F, Wiesenberger G, Adam G, Turgeon BG (2015) Individual and combined roles of malonichrome, ferricrocin, and TAFC siderophores in Fusarium graminearum pathogenic and sexual development. Front Microbiol 5: 759

Ponts N (2015) Mycotoxins are a component of Fusarium graminearum stress-response system. Front Microbiol 6: 1234

Pusztahelyi T, Holb I, Pócsi I (2015) Secondary metabolites in fungus-plant interactions. Front Plant Sci 6: 573

Robinson MD, McCarthy DJ, Smyth GK (2010) edgeR: a Bioconductor package for differential expression analysis of digital gene expression data. Bioinformatics 26: 139–140

Santhanam P, Thomma BPHJ (2012) Verticillium dahliae Sge1 differentially regulates expression of candidate effector genes. Mol Plant-Microbe Interact 26: 249–256

Sbaraini N, Andreis FC, Thompson CE, Guedes RLM, Junges Â, Campos T, Staats CC, Vainstein MH, Ribeiro de Vasconcelos AT, Schrank A (2017) Genome-wide analysis of secondary metabolite gene clusters in Ophiostoma ulmi and Ophiostoma novo-ulmi reveals a fujikurin-like gene cluster with a putative role in infection. Front Microbiol 8: 1063

Schrettl M, Bignell E, Kragl C, Sabiha Y, Loss O, Eisendle M, Wallner A, Arst Jr. HN, Haynes K, Haas H (2007) Distinct roles for intra-and extracellular siderophores during Aspergillus fumigatus infection. PLOS Pathog 3: e128

Shi-Kunne X, Faino L, van den Berg GCM, Thomma BPHJ, Seidl MF (2018) Evolution within the fungal genus Verticillium is characterized by chromosomal rearrangement and gene loss. Environ Microbiol 20: 1362–1373

Sieber CMK, Lee W, Wong P, Münsterkötter M, Mewes H-W, Schmeitzl C, Varga E, Berthiller F, Adam G, Güldener U (2014) The Fusarium graminearum genome reveals more secondary metabolite gene clusters and hints of horizontal gene transfer. PLOS One 9: e110311

Simms D, Cizdziel PE, Chomczynski P (1993) TRIzol: A new reagent for optimal single-step isolation of RNA. Focus 15: 532–535

Stamatakis A (2014) RAxML version 8: a tool for phylogenetic analysis and post-analysis of large phylogenies. Bioinformatics 30: 1312–1313

Tobiasen C, Aahman J, Ravnholt KS, Bjerrum MJ, Grell MN, Giese H (2007) Nonribosomal peptide synthetase (NPS) genes in Fusarium graminearum, F. culmorum and F. pseudograminearium and identification of NPS2 as the producer of ferricrocin. Curr Genet 51: 43–58

Trapnell C, Williams BA, Pertea G, Mortazavi A, Kwan G, van Baren MJ, Salzberg SL, Wold BJ, Pachter L (2010) Transcript assembly and quantification by RNA-Seq reveals unannotated transcripts and isoform switching during cell differentiation. Nat Biotechnol 28: 511

Tsuji G, Kenmochi Y, Takano Y, Sweigard J, Farrall L, Furusawa I, Horino O, Kubo Y (2002) Novel fungal transcriptional activators, Cmr1p of Colletotrichum lagenarium and Pig1p of Magnaporthe grisea, contain Cys2His2 zinc finger and Zn(II)2Cys6 binuclear cluster DNA-binding motifs and regulate transcription of melanin biosynthesis. Mol Microbiol 38: 940–954

Weber T, Blin K, Duddela S, Krug D, Kim HU, Bruccoleri R, Lee SY, Fischbach MA, Müller R, Wohlleben W, et al (2015) antiSMASH 3.0—a comprehensive resource for the genome mining of biosynthetic gene clusters. Nucleic Acids Res 43: W237–W243

Wiemann P, Keller NP (2014) Strategies for mining fungal natural products. J Ind Microbiol Biotechnol 41: 301–313

Xiong D, Wang Y, Tian L, Tian C (2016) MADS-Box transcription factor VdMcm1 regulates conidiation, microsclerotia formation, pathogenicity, and secondary metabolism of Verticillium dahliae. Front Microbiol 7: 1192

Yu X, Huo L, Liu H, Chen L, Wang Y, Zhu X (2015) Melanin is required for the formation of the multi-cellular conidia in the endophytic fungus Pestalotiopsis microspora. Microbiol Res 179: 1–11

Zhang D-D, Wang X-Y, Chen J-Y, Kong Z-Q, Gui Y-J, Li N-Y, Bao Y-M, Dai X-F (2016) Identification and characterization of a pathogenicity-related gene VdCYP1 from Verticillium dahliae. Sci Rep 6: 27979

